## Supplemental Figures and Tables for "Whole-organism 3D Quantitative Characterization of Zebrafish Melanin by Silver Deposition Micro-CT"

### Supplemental Material

**Figure S1**

**Figure S2**

**Figure S3**

**Supplementary Table 1**

**Supplementary Table 2**

**Supplementary Table 3**

**Supplementary Table 4**

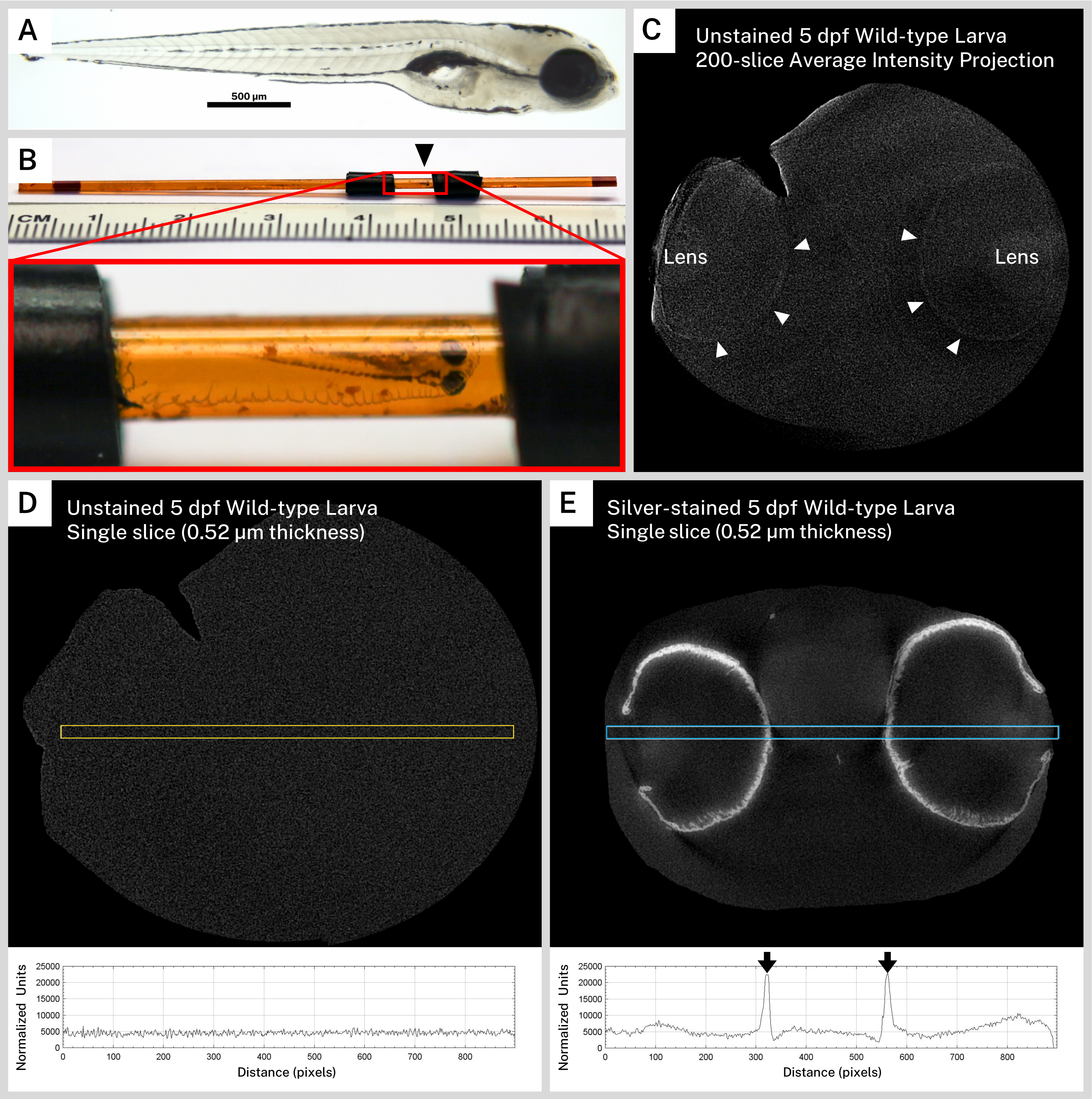


**Figure S1: Unstained zebrafish larvae do not exhibit pigment pattern intensity in micro-CT reconstructions.** A pigmented 5 dpf wild-type larva (**A**) was prepared for micro-CT imaging without silver staining. **B**) The sample (black arrowhead), embedded in resin within a polyimide tube, was surrounded by electrical tape to ensure proper positioning in the X-ray beam. **C**) A 200-slice average intensity projection of the normalized micro-CT reconstruction from this unstained sample shows faint attenuation in the lenses (L) and at the edges of the eyes (white arrowheads), indicating that the sample was scanned. A 900-pixel x 25-pixel selection (yellow box) transecting the eyes was used to calculate an intensity profile for a single reconstructed slice of the unstained fish (**D**) showing virtually no intensity peaks above background. A similar selection (blue box) through a single reconstructed slice of a silver-stained 5 dpf wild-type larva (**E**) demonstrates intense melanin staining in the intensity profile (black arrows).

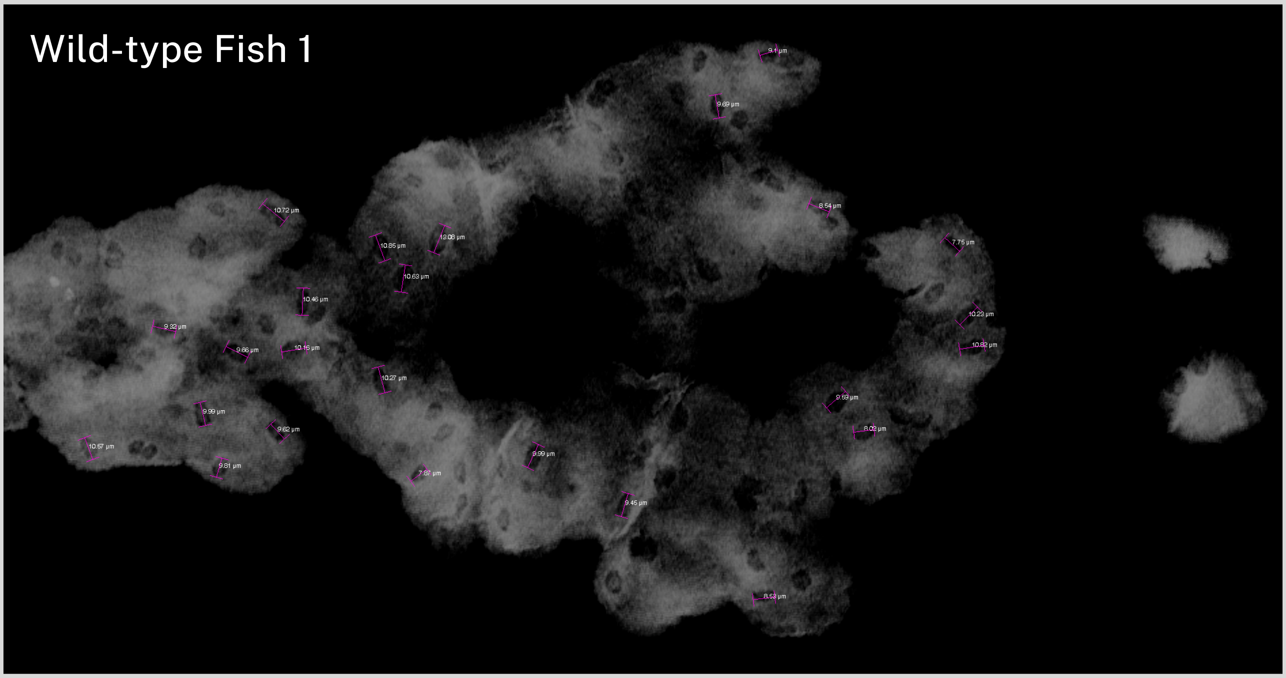
Wild-type Fish 1

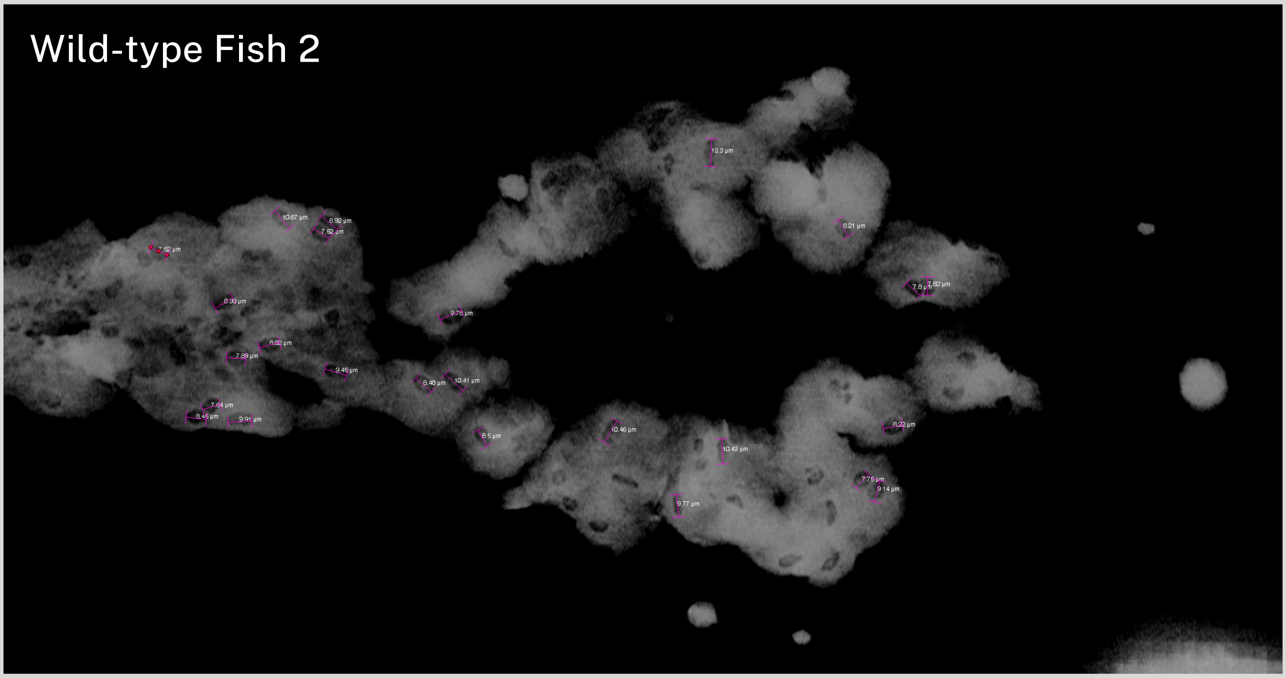
Wild-type Fish 2

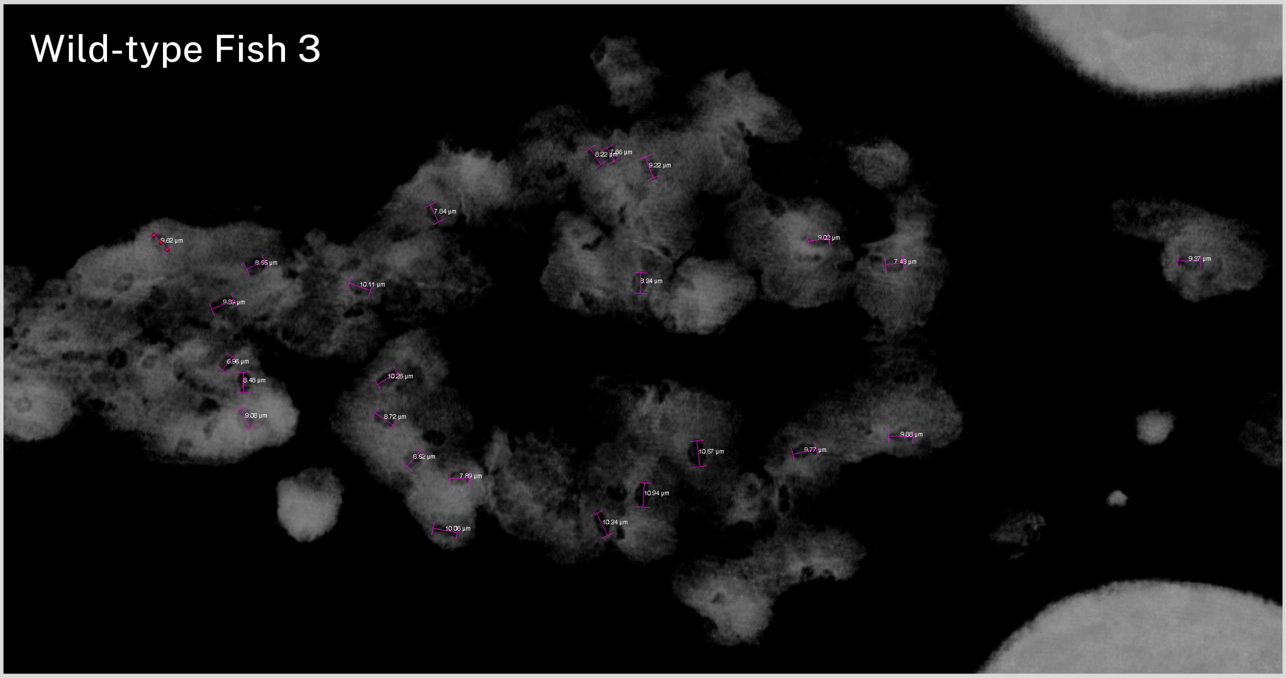
Wild-type Fish 3

**Figure S2: Measurements of ovoid transparencies in melanin staining.** Volume renderings of the top of the head for three 5 dpf wild-type zebrafish were examined for ovoid transparencies in the staining. For each fish, 25 random opacities were selected and manually measured on the long axis using the 3D line measurement tool in Avizo (pink lines). Average length (n=75) was 9.29 µm (S.D. = ±1.23 µm).

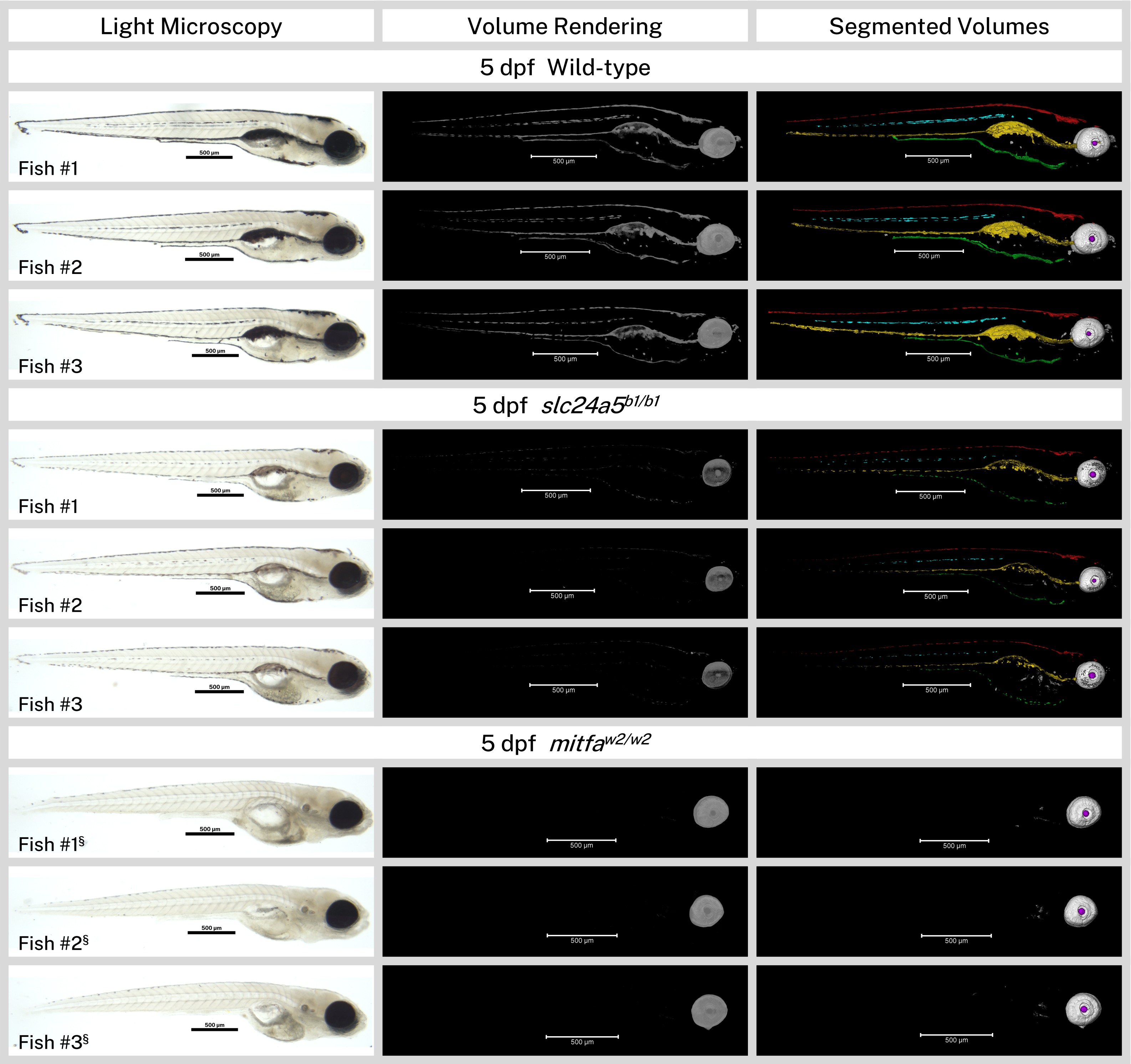


**Figure S3: Overview of all analyzed samples.** Light micrographs (before silver staining), micro-CT volume renderings, and segmented volumes of the wild-type, *slc24a5*$^{b1/b1}$, and *mitfa*$^{w2/w2}$ 5 dpf larvae (n=3 each) analyzed for this study. In the segmented volumes, Red = Dorsal stripe, Yellow = Ventral stripe, Green = Yolk Sac stripe, Cyan = Lateral stripes, White = RPE, Gray = other body melanin, Purple = Lens. Scale bars = 500 µm. § = head segment only analyzed by micro-CT and shown.

| Specimen | wt #1 | wt #2 | wt #3 | wt Average | S.D. |
|---|---|---|---|---|---|
| Pigment Region | Segmented Volume (µm³) | | | | |
| Total | 4,812,609 | 4,523,709 | 5,272,930 | 4,869,749 | ± 377,865 |
| RPE (right) | 1,203,302 | 1,022,566 | 1,276,510 | 1,167,459 | ± 130,711 |
| RPE (left) | 1,249,428 | 964,576 | 1,461,263 | 1,225,089 | ± 249,236 |
| Dorsal Stripe | 545,441 | 487,260 | 510,985 | 514,562 | ± 29,255 |
| Ventral Stripe | 1,406,879 | 1,556,401 | 1,621,673 | 1,528,318 | ± 110,117 |
| Yolk Sac Stripe | 228,825 | 363,388 | 205,678 | 265,964 | ± 85,162 |
| Lateral Stripe (right) | 52,441 | 41,027 | 67,341 | 53,603 | ± 13,195 |
| Lateral Stripe (left) | 59,082 | 29,626 | 61,595 | 50,101 | ± 17,776 |
| Other | 67,209 | 58,865 | 67,885 | 64,653 | ± 5024 |
| Pigment Region | Cumulative Sum of Intensity (normalized units) | | | | |
| Total | 505,334,686,000 | 470,665,376,000 | 472,890,158,300 | 482,963,406,767 | ± 19,406,004,611 |
| RPE (right) | 148,485,240,000 | 143,290,990,000 | 139,632,970,000 | 143,803,066,667 | ± 4,448,296,070 |
| RPE (left) | 174,322,350,000 | 146,697,760,000 | 157,178,020,000 | 159,399,376,667 | ± 13,945,619,823 |
| Dorsal Stripe | 48,810,881,000 | 40,159,785,000 | 40,131,047,000 | 43,033,904,333 | ± 5,003,029,185 |
| Ventral Stripe | 100,851,060,000 | 106,391,040,000 | 106,401,220,000 | 104,547,773,333 | ± 3,201,451,703 |
| Yolk Sac Stripe | 16,690,282,000 | 23,250,780,000 | 14,228,976,000 | 18,056,679,333 | ± 4,663,530,648 |
| Lateral Stripe (right) | 5,299,908,600 | 3,639,217,900 | 5,524,523,500 | 4,821,216,667 | ± 1,029,783,364 |
| Lateral Stripe (left) | 5,142,875,100 | 2,327,220,200 | 4,582,285,300 | 4,017,460,200 | ± 1,490,385,411 |
| Other | 5,732,089,300 | 4,908,582,900 | 5,211,116,500 | 5,283,929,567 | ± 416,553,724 |

**Supplementary Table 1: Quantification of Volume and Cumulative Sum of Intensity for Wild-type Samples for Segmented Pigment Regions.**

| Pigment Region | Segmented Volume Statistics | |
|---|---|---|
| | One-way ANOVA *F*-statistic | *P*-value |
| Total | F(2,6) = 178.43 | 0.000005 |
| RPE (right) | F(2,6) = 36.37 | 0.000442 |
| RPE (left) | F(2,6) = 21.77 | 0.002 |
| RPE (total) | F(2,6) = 29.64 | 0.001 |
| Dorsal Stripe | F(1,4) = 145.98 | 0.000269 |
| Ventral Stripe | F(1,4) = 376.02 | 0.000042 |
| Yolk Sac Stripe | F(1,4) = 22.04 | 0.009 |
| Lateral Stripe (right) | F(1,4) = 39.01 | 0.003 |
| Lateral Stripe (left) | F(1,4) = 17.48 | 0.014 |
| Other | F(2,6) = 52.86 | 0.000155 |
| Body (total) | F(2,6) = 939.89 | 0.000000032209 |
| Pigment Region | Cumulative Sum of Intensity Statistics | |
| | One-way ANOVA *F*-statistic | *P*-value |
| Total | F(2,6) = 185.76 | 0.000004 |
| RPE (right) | F(2,6) = 54.69 | 0.000141 |
| RPE (left) | F(2,6) = 70.40 | 0.000068 |
| RPE (total) | F(2,6) = 70.50 | 0.000068 |
| Dorsal Stripe | F(1,4) = 138.02 | 0.000300 |
| Ventral Stripe | F(1,4) = 1770.94 | 0.000002 |
| Yolk Sac Stripe | F(1,4) = 35.30 | 0.004 |
| Lateral Stripe (right) | F(1,4) = 56.79 | 0.002 |
| Lateral Stripe (left) | F(1,4) = 17.62 | 0.014 |
| Other | F(2,6) = 166.28 | 0.000006 |
| Body (total) | F(2,6) = 3690.79 | 0.00000000049447 |

**Supplemental Table 2: Statistical Analysis of Wild-Type, *slc24a5*$^{b1/b1}$, and *mitfa*$^{w2/w2}$ Samples for Segmented Pigment Regions.** F statistics for each comparison are reported as F(degrees of freedom between groups, degrees of freedom within groups) = F statistic. P-values <0.05 indicate a significant difference between groups. While the melanin stripes were compared between the wild-type and *golden* samples, total melanin, RPE melanin, and other melanin were compared between all three genotypes. For these regions, Tukey post-hoc tests were used to determine which groups differed significantly from wild-type.

| Specimen | *slc24a5*$^{b1/b1}$ #1 | *slc24a5*$^{b1/b1}$ #2 | *slc24a5*$^{b1/b1}$ #3 | *slc24a5*$^{b1/b1}$ Average | S.D. | % Change from wt | P-value |
|---|---|---|---|---|---|---|---|
| Pigment Region | Segmented Volume (μm$^3$) | | | | | | |
| Total | 1,140,958 | 1,324,056 | 1,141,011 | 1,202,008 | ± 105,696 | -75.3 | 0.000005* |
| RPE (right) | 344,581 | 459,837 | 449,497 | 417,972 | ± 63,768 | -64.2 | 0.000362* |
| RPE (left) | 408,323 | 415,649 | 405,002 | 409,658 | ± 5448 | -66.6 | 0.002* |
| Dorsal Stripe | 97,029 | 173,892 | 83,612 | 118,177 | ± 48,714 | -77.0 | 0.000269** |
| Ventral Stripe | 232,529 | 205,534 | 143,544 | 193,869 | ± 45,625 | -87.3 | 0.000042** |
| Yolk Sac Stripe | 39,712 | 39,309 | 20,702 | 33,241 | ± 10,861 | -87.5 | 0.009** |
| Lateral Stripe (right) | 4797 | 7575 | 4526 | 5632 | ± 1688 | -89.5 | 0.003** |
| Lateral Stripe (left) | 7155 | 9092 | 3991 | 6746 | ± 2575 | -86.5 | 0.014** |
| Other | 6833 | 13,167 | 30,136 | 16,712 | ± 12,049 | -74.2 | 0.001* |
| Pigment Region | Cumulative Sum of Intensity (normalized units) | | | | | | |
| Total | 75,413,111,000 | 85,132,595,270 | 74,976,816,050 | 78,507,507,440 | ± 5,741,640,000 | -83.7 | 0.000004* |
| RPE (right) | 24,331,246,000 | 31,653,982,000 | 31,072,209,000 | 29,019,145,667 | ± 4,070,247,833 | -79.8 | 0.000125* |
| RPE (left) | 29,773,920,000 | 31,726,170,000 | 28,370,035,000 | 29,956,708,333 | ± 1,685,517,492 | -81.2 | 0.000056* |
| Dorsal Stripe | 5,890,231,300 | 8,872,478,700 | 5,279,561,200 | 6,680,757,067 | ± 1,922,488,565 | -84.5 | 0.000300** |
| Ventral Stripe | 11,615,169,000 | 9,443,057,700 | 7,103,616,500 | 9,387,281,067 | ± 2,256,293,368 | -91.0 | 0.000002** |
| Yolk Sac Stripe | 2,589,991,900 | 1,841,591,900 | 1,261,461,400 | 1,897,681,733 | ± 666,038,944 | -89.5 | 0.004** |
| Lateral Stripe (right) | 310,585,660 | 407,258,620 | 274,772,610 | 330,872,297 | ± 68,533,178 | -93.1 | 0.002** |
| Lateral Stripe (left) | 481,020,900 | 452,190,750 | 243,505,540 | 392,239,063 | ± 129,611,111 | -90.2 | 0.014** |
| Other | 420,946,240 | 735,865,600 | 1,371,654,800 | 842,822,213 | ± 484,294,827 | -84.0 | 0.000015* |

**Supplementary Table 3: Quantification of Volume and Cumulative Sum of Intensity for *slc24a5*$^{b1/b1}$ Samples for Segmented Pigment Regions.** *p-values determined by Tukey post hoc test following one-way ANOVA. **p-values determined directly by one-way ANOVA. P-values were considered significant <0.05.

| Specimen | $mitfa^{w2/w2}$ #1§ | $mitfa^{w2/w2}$ #2§ | $mitfa^{w2/w2}$ #3§ | $mitfa^{w2/w2}$ Average | S.D. | % Change from wt | P-value |
|---|---|---|---|---|---|---|---|
| Pigment Region | Segmented Volume (μm³) | | | | | | |
| Total | 1,644,237 | 2,028,464 | 1,719,373 | 1,797,358 | ± 203,639 | -63.1 | 0.000015* |
| RPE (right) | 798,946 | 1,000,899 | 792,820 | 864,222 | ± 118,406 | -26.0 | 0.032* |
| RPE (left) | 841,073 | 1,022,548 | 921,264 | 928,295 | ± 90,942 | -24.2 | 0.120* |
| Other | 4218 | 5017 | 5289 | 4841 | ± 557 | -92.5 | 0.000169* |
| Pigment Region | Cumulative Sum of Intensity (normalized units) | | | | | | |
| Total | 196,770,044,220 | 263,788,449,540 | 191,002,155,630 | 217,186,883,130 | ± 40,461,050,760 | -55.0 | 0.000040* |
| RPE (right) | 101,128,420,000 | 132,800,940,000 | 88,039,498,000 | 107,322,952,667 | ± 23,014,687,673 | -25.4 | 0.040* |
| RPE (left) | 95,401,927,000 | 130,717,900,000 | 102,685,520,000 | 109,601,782,333 | ± 18,646,192,660 | -31.2 | 0.010* |
| Other | 239,697,220 | 269,609,540 | 277,137,630 | 262,148,130 | ± 19,804,054 | -95.0 | 0.000007* |

**Supplementary Table 4: Quantification of Volume and Cumulative Sum of Intensity for $mitfa^{w2/w2}$ Samples for Segmented Pigment Regions.** *p-values determined by Tukey post hoc test following one-way ANOVA. § = head segment only analyzed. P-values were considered significant <0.05.
